# A Quantitative Tractography Study into the Connectivity, Segmentation and Laterality of the Human Inferior Longitudinal Fasciculus

**DOI:** 10.1101/291625

**Authors:** Sandip S Panesar, Yeh Fang-Cheng, Timothée Jacquesson, William Hula, C Fernandez-Miranda Juan

**Affiliations:** Department of Neurological Surgery, University of Pittsburgh, Pittsburgh, United States of America; Department of Bioengineering, University of Pittsburgh, Pittsburgh, United States of America; Veterans Affairs Pittsburgh Healthcare System, Pittsburgh, United States of America

**Author notes:** **Corresponding Author** Juan C Fernandez-Miranda MD, Department of Neurological Surgery, University of Pittsburgh Medical Center, 200 Lothrop Street, Pittsburgh, 15213, Pennsylvania, United States of America.

## Abstract

The human inferior longitudinal fasciculus (ILF) is a ventral, temporo-occipital association tract. Though described in early neuroanatomical works, its existence was later questioned. Application of *in vivo* tractography to the neuroanatomical study of the ILF has generally confirmed its existence, however consensus is lacking regarding its subdivision, laterality and connectivity. Further, there is a paucity of detailed neuroanatomic data pertaining to the exact anatomy of the ILF. Generalized Q-Sampling imaging (GQI) is a non-tensor tractographic modality permitting high resolution imaging of white-matter structures. As it is a non-tensor modality, it permits visualization of crossing fibers and accurate delineation of close-proximity fiber-systems. We applied deterministic GQI tractography to data from 30 healthy subjects and a large-volume diffusion atlas, to delineate ILF anatomy. Post-mortem white matter dissection was also carried out in a cadaveric specimen for further validation. The ILF was found in all 60 hemispheres. At its occipital extremity, it demonstrated a trifurcated termination pattern which was used to separate the ILF into 3 distinct sub-fascicles: *Dorsolateral, ventrolateral and ventromedial.* These divisions were consistent across the subject set and within the atlas. We applied quantitative techniques to study connectivity strength of the ILF at its anterior and posterior extremities. Overall, the 3 sub-fascicles, and the whole ILF, demonstrated strong leftward-lateralized connectivity patterns. Leftward-lateralization was also found for ILF volumes across the subject set. Due to connective and volumetric leftward-dominance and ventral location, we postulate the ILFs role in the semantic system. Further, our results are in agreement with functional and lesion-based postulations pertaining to the ILFs role in facial recognition.

## 1. Introduction

The inferior longitudinal fasciculus (ILF) is a human association tract situated in the ventral temporal lobe. Historically, it was described as connecting the superior, middle, inferior and fusiform gyri, to the lingual, cuneate, lateral-occipital and occipito-polar cortices. Early descriptions came from post-mortem white matter dissection (Crosby, 1962; Dejerine and Dejerine-Klumpke, 1895; Gloor, 1997). More recently, radioisotopic tracer studies in non-human primates questioned the existence of a robust temporo-occipital fasciculus, proposing instead a series of cortico-cortical U-fibers, travelling along the antero-posterior temporal distance (Catani et al., 2003; Tusa and Ungerleider, 1985). The introduction of tractography somewhat reconciled this controversy: Early diffusion tensor imaging (DTI) studies confirmed the existence of the human ILF (Catani et al., 2002, 2003; Fernández-Miranda et al., 2008; Jellison et al., 2004; Kier et al., 2004; Urbanski et al., 2008; Wakana et al., 2004). Catani et al. (2002, 2003) described a bi-component structure, with a direct temporo-occipital sub-tract, and an indirect series of cortico-cortical U-fibers. The direct subfascicle demonstrated a tri-pronged posterior, and bi-pronged anterior connectivity profile. This connective and structural description remained largely unchanged until a recent dissection and DTI series by Latini et al. (2015, 2017). The authors proposed a rightward-lateralized, dorsal-ventral ILF arrangement derived from its posterior connectivity profile: dorsally, distinct ILF subfasciculi originated from the cuneate and lateral occipital lobes, while ventrally, subfasciculi originated from the fusiform and lingual gyri (Latini, 2015; Latini et al., 2017).

Prior to the introduction of functional neuroimaging, data pertaining to ILF function was derived primarily from lesional data. Visual agnosia and amnesia were attributed to bilateral damage to ventral temporal white matter. Prosapagnosia (Bauer, 1984; Benson, 1974; Grossi et al., 2014; Michel et al., 1989) and visual hypo-emotionality (Fischer et al., 2016; Habib, 1986; Takeuchi et al., 2013) were linked to right-sided ventral temporal lesions. Further insights have come from both functional neuroimaging (e.g. fMRI) and intraoperative electrical stimulation (IES) (Duffau et al., 2005; Mandonnet et al., 2007). fMRI studies demonstrated semantic activations within temporo-polar, anterior temporal, basal occipito-temporal and occipital lobes corresponding to dominant-hemisphere ILF trajectory (Saur et al., 2008, 2010). An IES study by Mandonnet et al. (2007) proposed the ILF’s role in semantic functionality, consistent with the ‘dorsal-ventral’ model of language organization (Hickok and Poeppel, 2015).

Tractography permits functional insights by studying cortical-connectivity patterns of white matter tracts. It also allows calculation of volumetric, lateralization and subdivision patterns of white matter systems. It therefore surpasses post-mortem dissection in terms of ability and accuracy (Fernandez-Miranda et al., 2012). DTI is unable to delineate crossing fibers and demonstrate cortical connectivity accurately (Farquharson et al., 2013; Fernandez-Miranda et al., 2012). Generalized q-sampling imaging (GQI) dispenses with the diffusion tensor, allowing crossing fibers at close proximity to be tracked at high-resolution (Yeh et al., 2010, 2013). Our group has pioneered the application of GQI tractography to neuroanatomical (Fernández-Miranda et al., 2015; Meola et al., 2016; Panesar et al., 2017; Wang et al., 2013, 2016; Yoshino et al., 2016) and surgical studies (Fernandez-Miranda et al., 2012). Further, our use of large-volume GQI-derived atlases (Fernández-Miranda et al., 2015; Panesar et al., 2017; Wang et al., 2013; Yeh and Tseng, 2011) provides a model of *‘*average*’* white matter anatomy for validation. With these considerations in mind, we set out to study the subdivision, asymmetry and connectivity of the human ILF using GQI tractography using quantitative connectometric methods, along with diffusion atlas and dissection validation.

## 2. Methods

### 2.1 Participants

We conducted a subject-specific deterministic fiber tractography study in 30 right-handed, nurologically-healthy male and female subjects, age range 23-35. The data were from the Human Connectome Project (HCP) online database (WU-Minn Consortium (Principal Investigators: David van Essen and Kamil Ugurbil; 1U54MH091657) funded by the 16 NIH institutes and centers that support the NIH Blueprint for Neuroscience Research and by the McDonnell Center for Systems Neuroscience at Washington University. Likewise, data from 842 individual HCP subjects were utilized to compile the averaged diffusion atlas.

### 2.2 Image Acquisition and Reconstruction

The HCP diffusion data for individual subjects were acquired using a Siemens 3T Skyra system, with a 32-channel head coil (Siemans Medical, Erlangen, Germany). A multishell diffusion scheme was used, and the *b* values were 1000, 2000, and 3000 s/mm^2^. The number of diffusion sampling directions were 90, 90 and 90, respectively. The in-plane resolution and slice thickness were both 1.25mm (TR = 5500 ms, TE = 89 ms, resolution = 1.25mm x 1.25mm, FoV = 210mm x 180mm, matrix = 144 x 168). The DSI data were reconstructed using the generalized q-sampling imaging approach (Yeh et al., 2010) using a diffusion distance ratio of 1.2 as recommended by the original study.

### 2.3 HCP 842 Atlas

A total of 842 participants from the HCP database were used to construct the atlas. The image acquisition parameters are identical to those described previously. The diffusion data were reconstructed and warped to the Montreal Neurological Institute (MNI) space using *q*-space diffeomorphic reconstruction (Yeh and Tseng, 2011) with a diffusion sampling length ratio of 1.25 and the output resolution was 1 mm. The group average atlas was then constructed by averaging the reconstructed data of the 842 individual subjects within the MNI space.

### 2.4 Fiber Tracking and Analysis

We performed deterministic fiber tracking using DSI Studio software, which utilizes a generalized streamline fiber tracking method (Yeh, Verstynen, Wang, Fernandez-Miranda, & Tseng, 2013). Parameters selected for fiber tracking included a step size of 0.2 mm, a minimum fiber length of 20mm and a turning angle threshold of 60°. For progression locations containing >1 fiber orientation, fiber orientation most congruent with the incoming direction and turning angle <60° was selected to determine subsequent moving direction. Each progressive voxels’ moving directional estimate was weighted by 20% of the previous voxels incoming direction and by 80% of its nearest fiber orientation. This sequence was repeated to create fiber tracts. Termination of the tracking algorithm occurred when the quantitative anisotropy (QA) (Yeh et al., 2013) dropped below a subject-specific value: when fiber tract continuity no longer met the progression criteria, or when 100,000 tracts were generated. We pre-selected QA termination threshold, between 0.02-0.08, by analyzing the number of false continuities generated within each subjects’ dataset and chose the compromise value that allowed optimal anatomical detail with minimal noise. Likewise, we selected a smoothing parameter of 50% for the same reason stated previously.

### 2.5 Region of Interest Placement and Fiber Selection

Unlike our previous studies, and in an effort to replicate published methodology (Catani et al., 2003; Latini et al., 2017), we chose a two region of interest (ROl) approach rather than an atlas-based seeding approach. This method was chosen to minimize *a priori* selection of feed-backward or feed-forward tracts. A spherical ROI was positioned within the white matter of the anterior temporal lobe. A second, rectangular ROI was placed in the coronal plane at the temporo-occipital junction. To avoid commissural fibers and false continuities, a rectangular region of avoidance was placed across the sagittal hemispheric midline. Once 100,000 fiber tracts were generated, we manually removed fibers passing through the ventral external capsule (i.e. inferior fronto-occipital (IFOF), uncinate (UF) fasciculi) and fibers resembling the arcuate fasciculus (AF), superior longitudinal fasciculus (SLF) and middle longitudinal fasciculus (MdLF) **(Figure 1A).**

**Figure 1.**
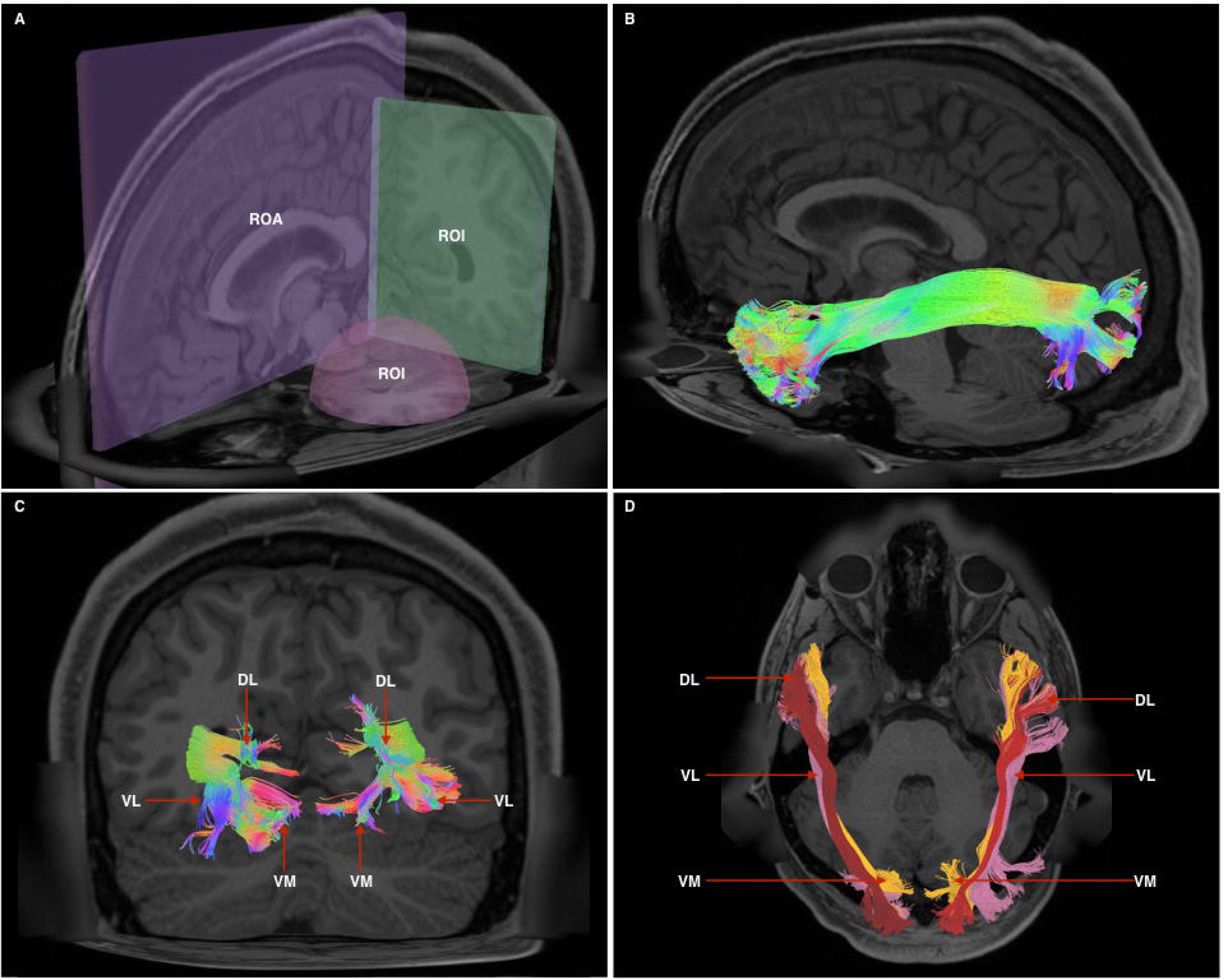
1A - Cutaway view showing all 3 radiological planes (axial, sagittal, coronal) with ROls and ROAs used. Spherical ROl is placed within the anterior temporal lobe white matter, whilst the rectangular ROl is placed in a coronal plane at the approximate position of the temporo-occipital junction. The ROA was placed in the mid-sagittal plane to exclude any fibers crossing the midline. 1B - Generated whole, undivided ILF bundle after manual removal of spurious fibers belonging to other white matter systems and prior to separation. Color assignment is directional. 1C - A posterior-coronal view demonstrating the posterior terminations of both left and right ILF bundles. Visible clearly is the trifurcated arrangement of each whole-ILF bundle which were subsequently used as geographical landmarks to separate the ILF further. DL - dorsolateral, VL - ventrolateral, VM - ventromedial. 1D - A superior-axial view demonstrating left and right ILF bundles following separation of the ILF into its respective subdivisions. The sub-fascicles have been individually colored. DL - dorsolateral (red), VL-ventrolateral (pink), VM - ventromedial (orange).

### 2.6 Quantitative Connectivity Analysis and Connectogram Creation

We used a quantitative method to define cortical terminations. Each sub-fascicle was merged with its hemispheric counterpart (i.e. left and right). The *‘*connectivity matrix*’* function in DSI studio was used to generate matrices representing the number of fibers terminating within regions of a specially modified, per-subject aligned version of the Automated Anatomical Labelling (AAL) atlas. Three connectivity matrices were generated per subject, to give a total of 90 connectivity matrices corresponding to *dorsolateral, ventrolateral, ventromedial* sub-fascicles (see results) over the 30 individual subjects. These matrices were collated in Matlab and the number of fibers corresponding to each respective connection were divided by the total number of fibers for each fascicle, per subject. Values were then scaled over the range of 30 subjects to give a connection index (Cl) between 0-100, with 0 representing no connectivity, and 100 representing strongest relative connectivity to a particular atlas region. Index values were then used to generate connectograms for each sub-fascicle and for whole ILF bundles, using CIRCOS (http://mkweb.bcgsc.ca/tableviewer/visualize/). Connectograms provide a unique method of visualizing network topology by demonstrating weighted strength of connectivity between brain regions.

### 2.7 Volumetry and Lateralization

We calculated the number of voxels occupied by each fiber trajectory (streamlines) and the subsequent volume (in milliliters) of each gross ILF bundle and their manually separated subfascicles. The volumes of left and right merged ILFs and sub-fascicles across 30-subjects were each subjected to an independent samples *T*-test in SPSS (IBM Corporation, Armonk, New York) to calculate significance of mean hemispheric volumetry over the 30 subjects. We also calculated the lateralization index (LI) using the formula (((volume left - volume right) ÷ (volume left + volume right)) x 2) (Catani et al., 2007).

### 2.8 White Matter Dissection

Three human hemispheres were prepared for dissection. First, they were fixed in a 10% formalin solution for two months. After fixation, the arachnoid and superficial vessels were removed. The brains were subsequently frozen at -16^°^C for two weeks, as per the Klingler method (Ludwig and Klingler, 1956). Dissection commenced 24 hours after the specimens were thawed and proceeded in a step-wise, superficial-to-deep process, at the lateral and inferior surface of the temporal lobe. Dissection was achieved using wooden spatulas to remove successive layers of grey matter followed by white matter.

## 3 Results

### 3.1 Morphology and Subdivision of ILF Bundles

Bundles resembling the ILF were found in 60/60 hemispheres **(Figure 1B)** in our subject-specific tractographic analysis and bilaterally within the 842-subject template. Upon generating and extracting whole ILF bundles we observed a distinct posterior termination pattern across the individual subjects and in the template, which we used to further divide the ILFs **(Figure 1C).** Though the whole ILF bundles demonstrated a distinct posterior termination morphology, their anterior aspects did not. We chose to further subdivide ILFs according to their posterior arrangement, terming the subfascicles *‘dorsolateral,’ ‘ventrolateral’* and *‘ventromedial’* ILF, respectively **(Figure 1D, 6A-B).** The dorsolateral bundle was found in 60/60 hemispheres. The ventrolateral bundle was found in 28/30 left and 27/30 right hemispheres, respectively. The ventromedial bundle was found in 30/30 left and 29/30 right hemispheres. Left and right ILFs were successfully replicated on the 842-subject atlas and assumed identical morphology and subdivisions compared to those from the subject-specific study.

### 3.2 Quantitative ILF Connectivity

#### 3.2.1 Dorsolateral ILF Sub-Fascicles (Table 1.)

The left dorsolateral ILF bundle demonstrated strongest connectivity patterns between the superior occipital to superior temporal gyri (Cl: 100.00) and middle temporal gyri (Cl: 41.13). All other connections had a Cl <40. Right sided dorsolateral ILF connectivity was comparatively weaker versus the left: maximum connectivity was found between the cuneus and superior temporal gyrus (Cl: 25.99), with all other connections demonstrating Cl <25. Left dorsolateral ILF connectivity was spread over 9 occipito-temporal connections, while the right was spread over 7. The average Cl for the left dorsolateral ILF was 14.77 (SD = 26.13) for the left, while it was 10.45 (SD = 15.83) for the right (Not significant, *T =* 0.61, p = 0.547). Median Cl for the left dorsolateral ILF was 23.10 and was 5.90 on the right **(Figure 2).**

**Table 1.**
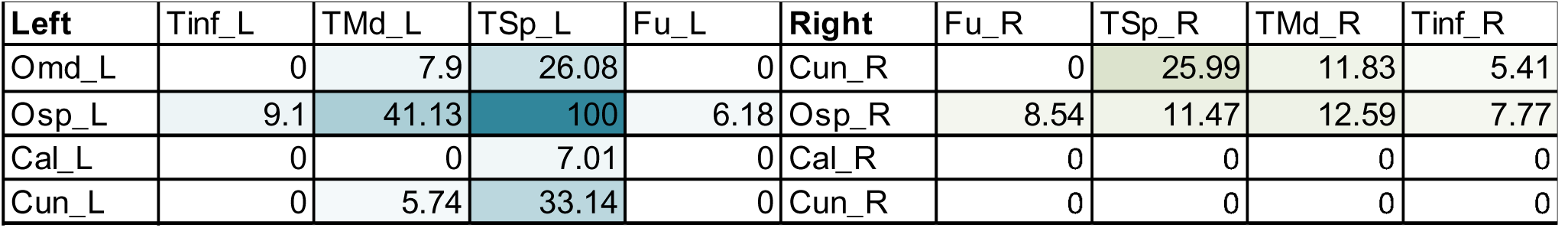
Table of connectivity indices for left (blue) and right (green) dorsolateral ILF sub-fascicles. Darker color represents stronger index.

| Left | Tinf_L | TMd_L | TSp_L | Fu_L | Right | Fu_R | TSp_R | TMd_R | Tinf_R |
| --- | --- | --- | --- | --- | --- | --- | --- | --- | --- |
| Omd_L | 0 | 7.9 | 26.08 | 0 | Cun_R | 0 | 25.99 | 11.83 | 5.41 |
| Osp_L | 9.1 | 41.13 | 100 | 6.18 | Osp_R | 8.54 | 11.47 | 12.59 | 7.77 |
| Cal_L | 0 | 0 | 7.01 | 0 | Cal_R | 0 | 0 | 0 | 0 |
| Cun_L | 0 | 5.74 | 33.14 | 0 | Cun_R | 0 | 0 | 0 | 0 |

#### 3.2.2 Ventrolateral ILF Sub-Fascicles (Table 2.)

The left ventrolateral ILF demonstrated strongest connectivity between lingual and middle temporal gyrus (Cl: 100.00), followed by lingual to superior temporal gyrus (Cl: 69.80), inferior occipital to inferior temporal gyrus (Cl: 68.69), middle occipital to superior temporal gyrus (Cl: 63.41) and inferior occipital to middle temporal gyrus (Cl: 57.88). All other left sided connections had a Cl <50. Versus the left, the right ventrolateral ILF demonstrated relatively weaker connectivity indices. Strongest right-sided CIs were lingual to middle temporal gyrus (Cl: 60.99), inferior occipital to inferior temporal gyrus (Cl: 51.22) and middle temporal gyrus (Cl: 39.18). All other connections had Cl <20. Both left and right ventrolateral ILFs demonstrated spread over 17 connections each. The average Cl for the left ventrolateral ILF was 36.45 (SD = 28.07) for the left, while it was 15.57 (SD = 17.70) for the right (Significant, *T =* 2.90, p = 0.006). Median Cl for the left ventrolateral ILF was 36.59 and was 10.03 for the left **(Figure 3).**

**Figures 2-5 Legend.**
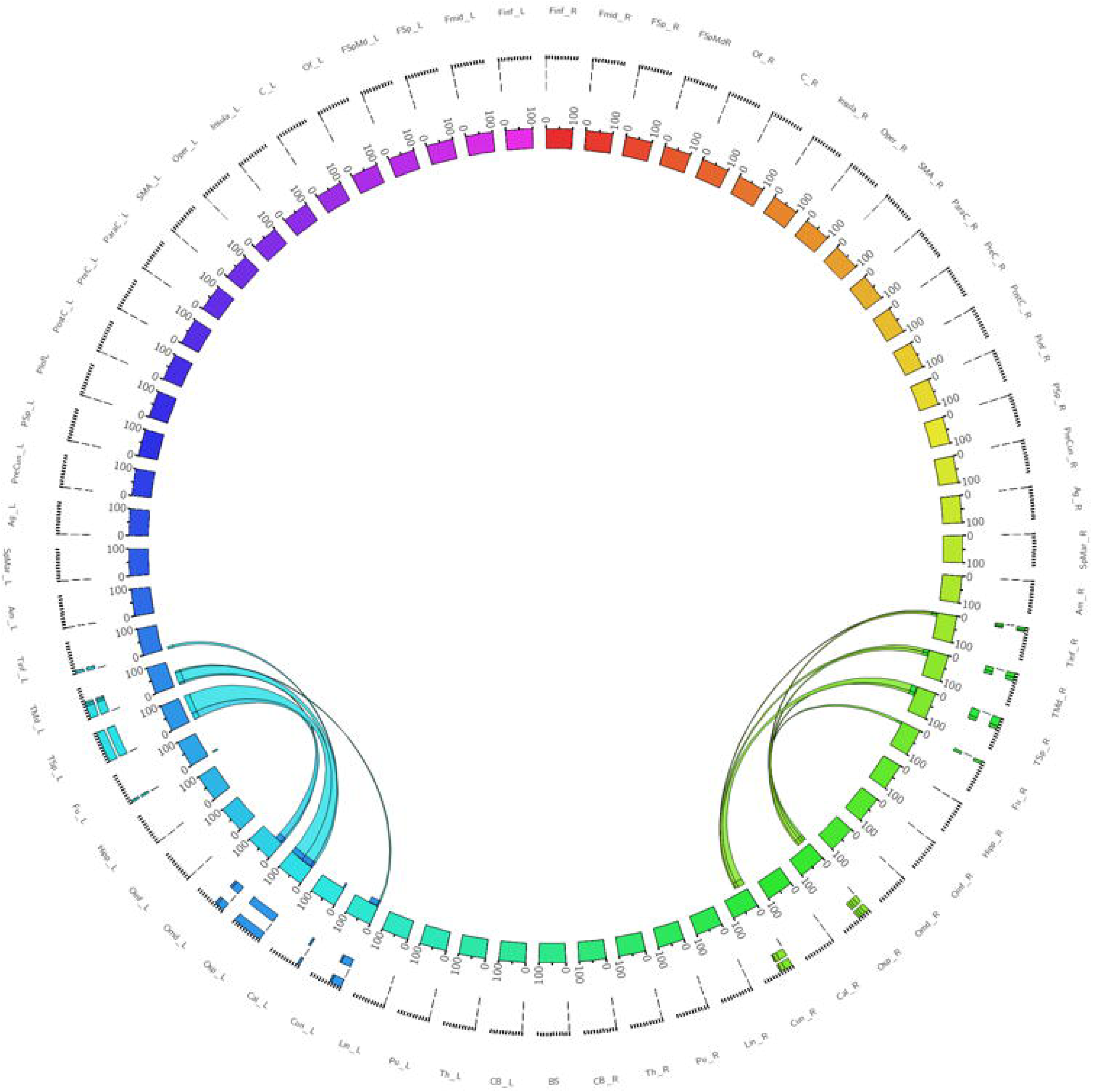
(NB: Only pertinent AAL atlas regions are listed) Tlnf - Inferior temporal gyrus, TMd - Middle temporal gyrus, TSp-Superior temporal gyrus, Fu - Fusiform gyrus, Olnf-inferior occipital gyrus, OMd - middle occipital gyrus, OSp - superior occipital gyrus, Cal - calcarine gyrus, Cun - cuneus, Lin - lingual gyrus. Left and Right hemispheric connections are demonstrated with a suffix _L or _R Figure 2 - A connectogram representing bilateral connectivity patterns of the dorsolateral ILF subfascicle. Atlas regions are listed around the circumference of the connectogram.

**Table 2.** Table of connectivity indices for left (blue) and right (green) ventrolateral ILF sub-fascicles. Darker color represents stronger index.

| Left | Tinf_L | TMd_L | TSp_L | Fu_L | Right | Hipp_R | Fu_R | TSp_R | TMd_R | Tinf_R |
| --- | --- | --- | --- | --- | --- | --- | --- | --- | --- | --- |
| Fu_L | 25.21 | 27.47 | 6 | 0 | Lin_R | 0 | 10.03 | 37.47 | 60.99 | 37.24 |
| Oinf_L | 68.69 | 57.88 | 45.42 | 7.11 | Cal_R | 0 | 0 | 9.48 | 16.2 | 8.77 |
| Omd_L | 37.54 | 37.74 | 63.41 | 0 | Omd_R | 0 | 0 | 14.43 | 6.23 | 0 |
| Cal_L | 44.32 | 35.63 | 34.5 | 0 | Oinf_R | 5.47 | 13.79 | 35.38 | 39.18 | 51.22 |
| Lin_L | 61.13 | 100 | 69.8 | 7.22 | Fu_R | 0 | 0 | 10.24 | 15.05 | 18.15 |

**Figure 3.**
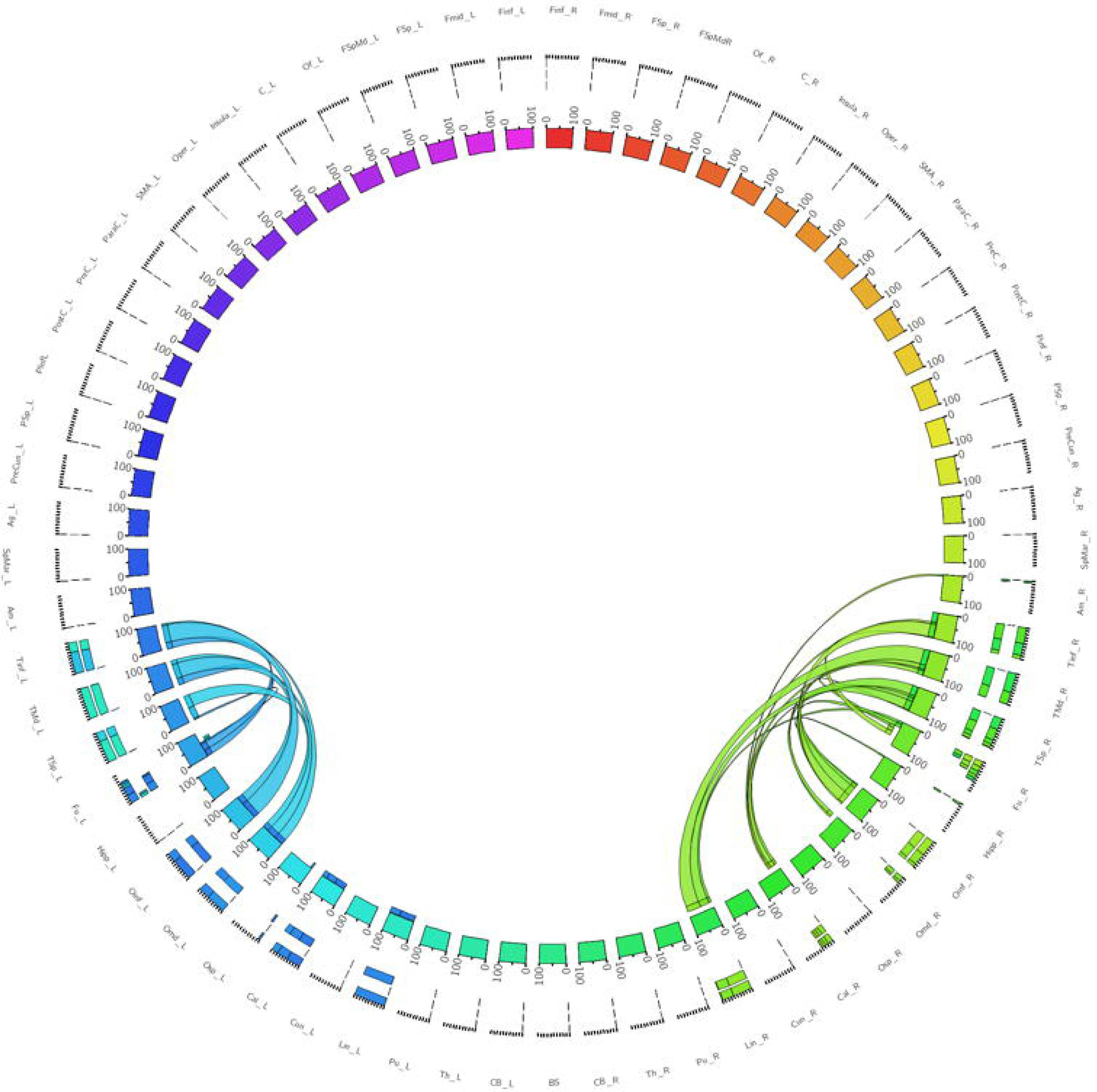
A connectogram representing bilateral connectivity patterns of the ventrolateral ILF subfascicle. Atlas regions are listed around the circumference of the connectogram.

#### 3.2.3 Ventromedial ILF Sub-Fascicles (Table 3.)

The left ventromedial ILF demonstrated strongest connectivity between the lingual and middle temporal gyrus (Cl: 81.58), calcarine to middle temporal gyrus (Cl: 79.07) and calcarine to superior temporal gyrus (Cl: 71.35). All other connections had Cl <70. The right ventromedial ILF demonstrated the strongest connectivity between lingual and middle temporal gyrus (Cl: 100.00) followed by lingual to inferior <20. The left ventromedial ILF connectivity was spread over 17 occipito-temporal connections, while the right was spread over 13. The average Cl for the left ventromedial ILF was 22.88 (SD = 28.15) or the left, while it was 14.44 (SD = 24.09) for the right (Not significant, *T =* 1.07, p = 0.290). Median Cl for the left ventrolateral ILF was 11.20 and was 6.50 for the left **(Figure 4).**

**Table 3.** Table of connectivity indices for left (blue) and right (green) ventromedial ILF sub-fascicles. Darker color represents stronger index.

| Left | Tinf_L | TMd_L | TSp_L | Fu_L | Right | Fu_R | TSp_R | TMd_R | Tinf_R | Am_R |
| --- | --- | --- | --- | --- | --- | --- | --- | --- | --- | --- |
| Oinf_L | 0 | 16.37 | 19.26 | 0 | Lin_R | 14.45 | 37.59 | 100.00 | 51.32 | 6.63 |
| Omd_L | 7.35 | 21.95 | 21.03 | 0 | Cal_R | 5.26 | 12.56 | 12.96 | 13.76 | 6.74 |
| Osp_L | 5.95 | 10.86 | 0 | 0 | Oinf_R | 0.00 | 15.20 | 6.37 | 0.00 | 0.00 |
| Cal_L | 55.99 | 79.07 | 71.35 | 6.5 | Fu_R | 0.00 | 0.00 | 6.13 | 0.00 | 0.00 |
| Cun_L | 12.4 | 11.53 | 0 | 0 | Cun_R | 0.00 | 0.00 | 0.00 | 24.09 | 14.45 |
| Lin_L | 56.64 | 81.58 | 65.72 | 5.52 | Lin_R | 0.00 | 0.00 | 0.00 | 0.00 | 0.00 |

**Figure 4.**
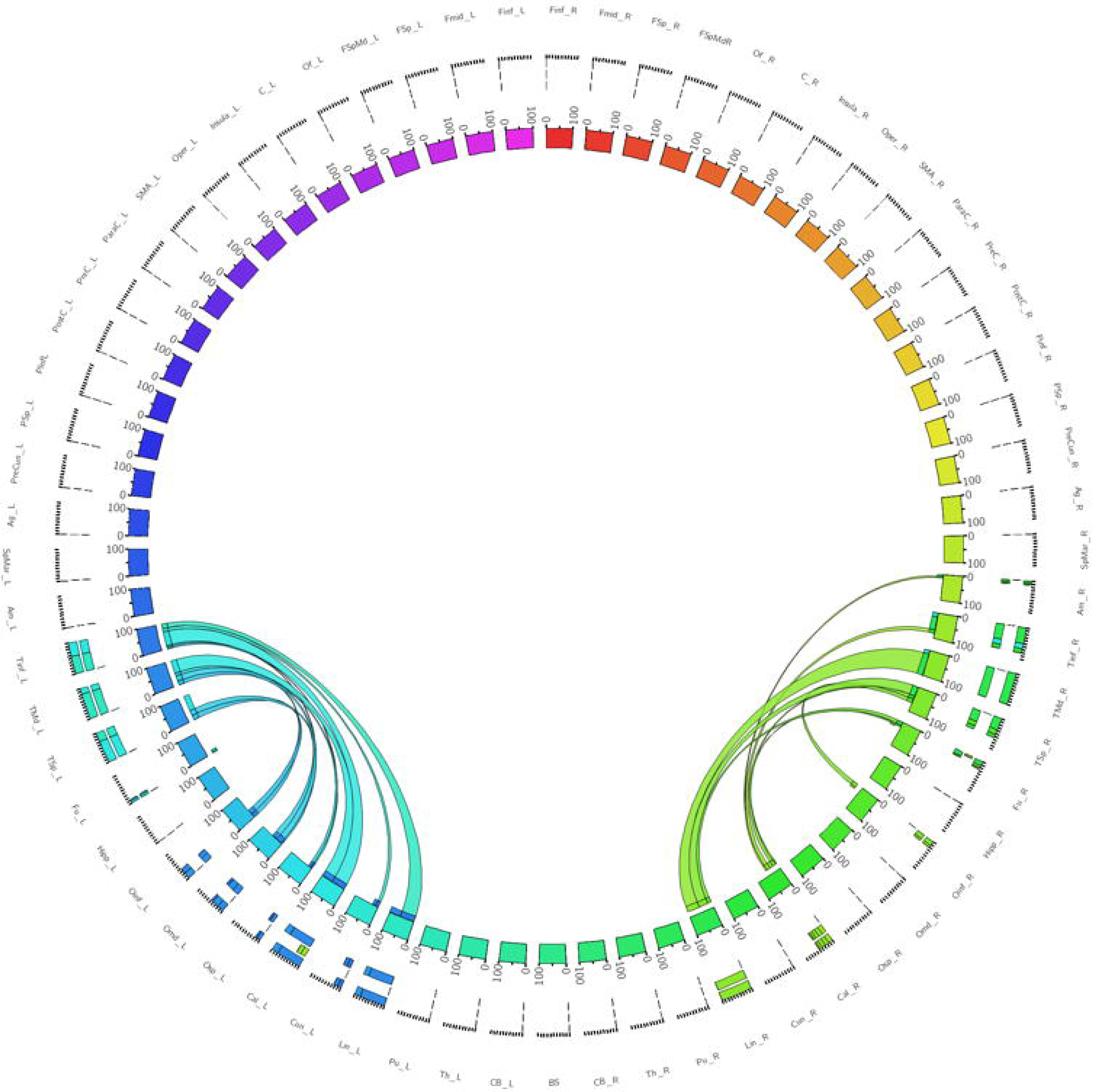
A connectogram representing bilateral connectivity patterns of the ventromedial ILF subfascicle. Atlas regions are listed around the circumference of the connectogram.

#### 3.2.4 Whole ILF Bundles (Table 4.)

When taken as whole, merged bundles, both left and right ILFs each demonstrated 22 patterns of occipito-temporal connectivity, respectively. The left ILF, however demonstrated stronger connectivity indices versus the right. On the left, strongest connectivity was demonstrated between lingual to middle temporal gyrus (Cl: 100.00), superior occipital to superior temporal gyrus (Cl: 77.81), lingual to superior temporal gyrus (Cl: 77.67), calcarine to middle temporal gyrus (Cl: 69.94) and calcarine to superior temporal gyri (Cl: 63.18). All other connections had Cl <60. The strongest connections on the right side were between lingual and middle (Cl: 77.54), inferior (Cl: 41.05) and superior (Cl: 40.70) temporal gyri. All other connections had a Cl <40. The average Cl for the whole ILF was 29.50 (SD = 29.29) for the left, while it was 10.60 (SD = 15.83) for the right (Significant, *T =* 3.07, p = 0.004). Median Cl for the left ventrolateral ILF was 23.10 and was 5.90 for the right **(Figure 5, Figure 6C-D, Figure 7A).**

**Table 4.**
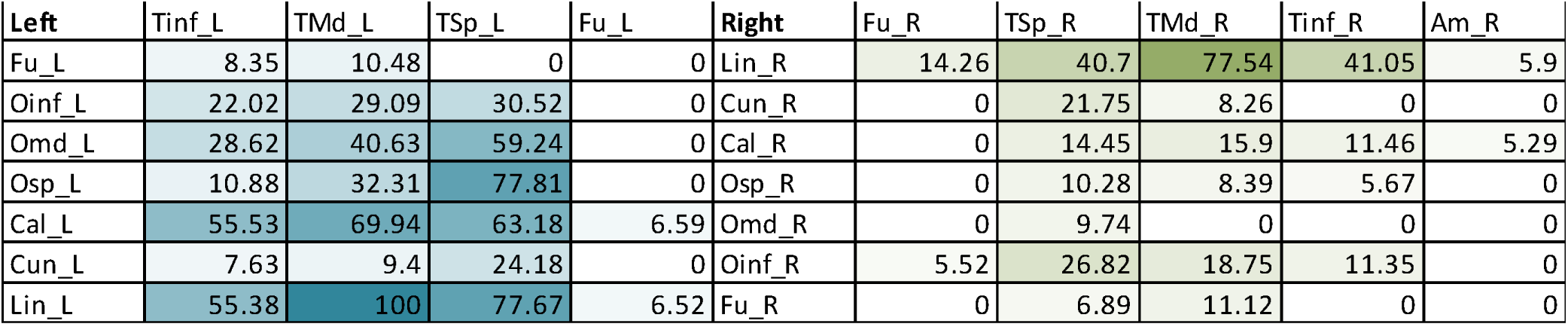
Table of connectivity indices for left (blue) and right (green) whole ILFs. Darker color represents stronger index.

**Figure 5.**
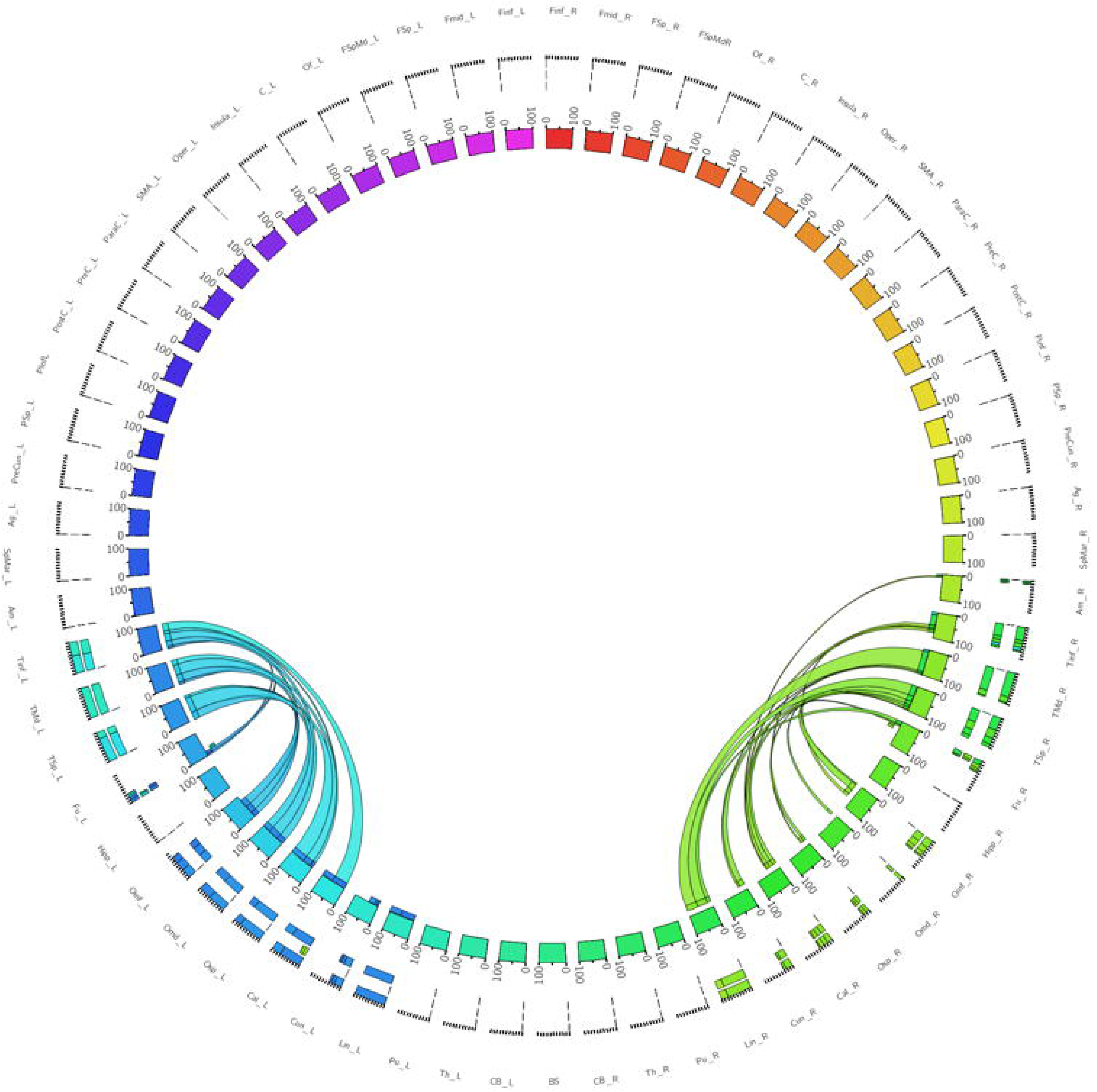
A connectogram representing bilateral connectivity patterns of the whole, unseparated ILF. Atlas regions are listed around the circumference of the connectogram.

**Figure 6.**
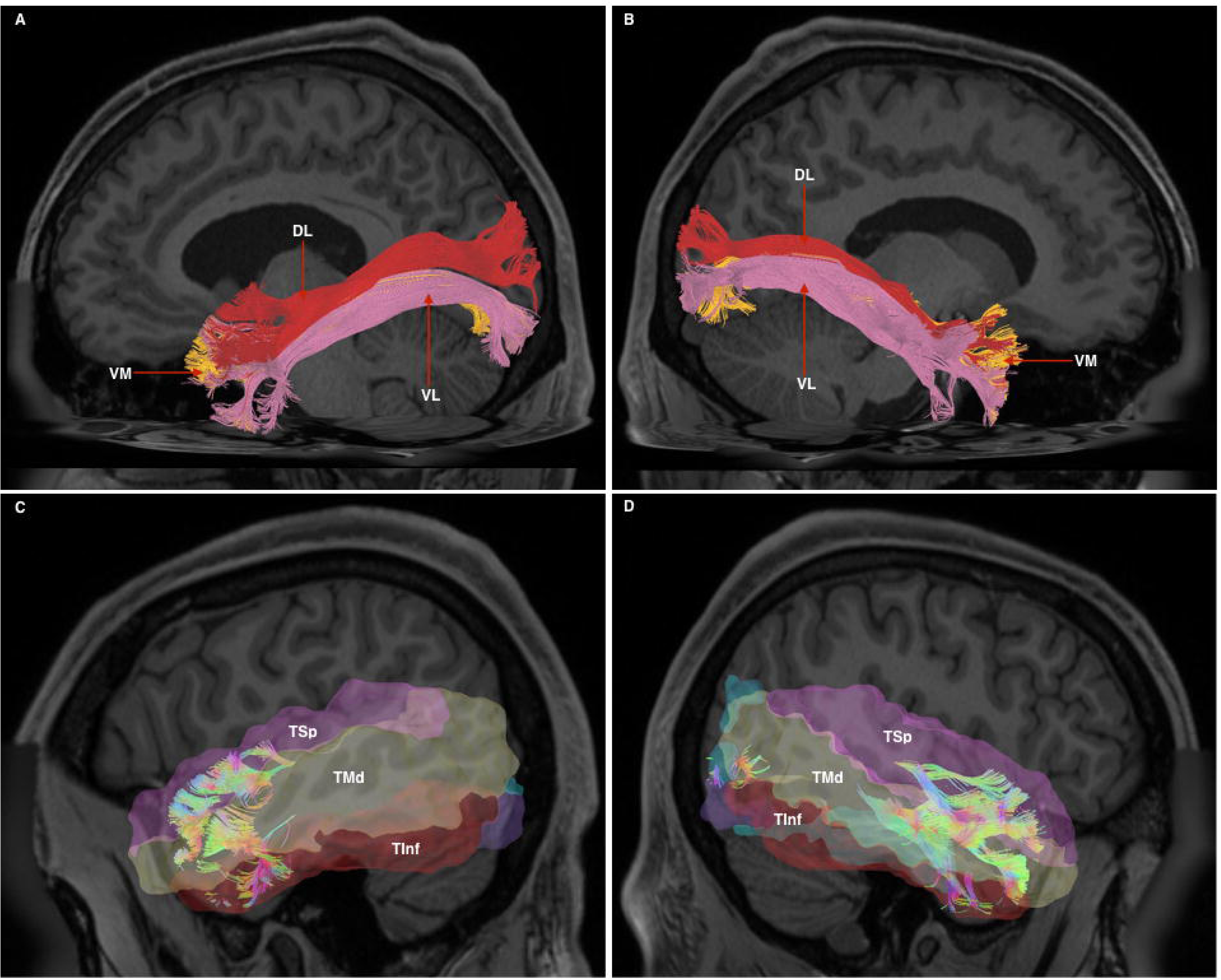
6A - Left-hemispheric, sagittal-view representation of a subdivided ILF. DL - dorsolateral (red), VM - ventromedial (orange), VL-ventrolateral (red). 6B - Right-hemispheric, sagittal-view representation of a subdivided ILF. DL - dorsolateral (red), VM - ventromedial (orange), VL-ventrolateral (red). 6C - Left-hemispheric, sagittal-view representation of anterior connectivity pattern of the whole, undivided ILF. Atlas regions corresponding to the superior (TSp), middle (TMd) and inferior (Tlnf) temporal gyri have been placed to demonstrate connectivity. 6D - Right-hemispheric, sagittal-view representation of anterior connectivity pattern of the whole, undivided ILF. Atlas regions corresponding to the superior (TSp), middle (TMd) and inferior (Tlnf) temporal gyri have been placed to demonstrate connectivity.

**Figure 7.**
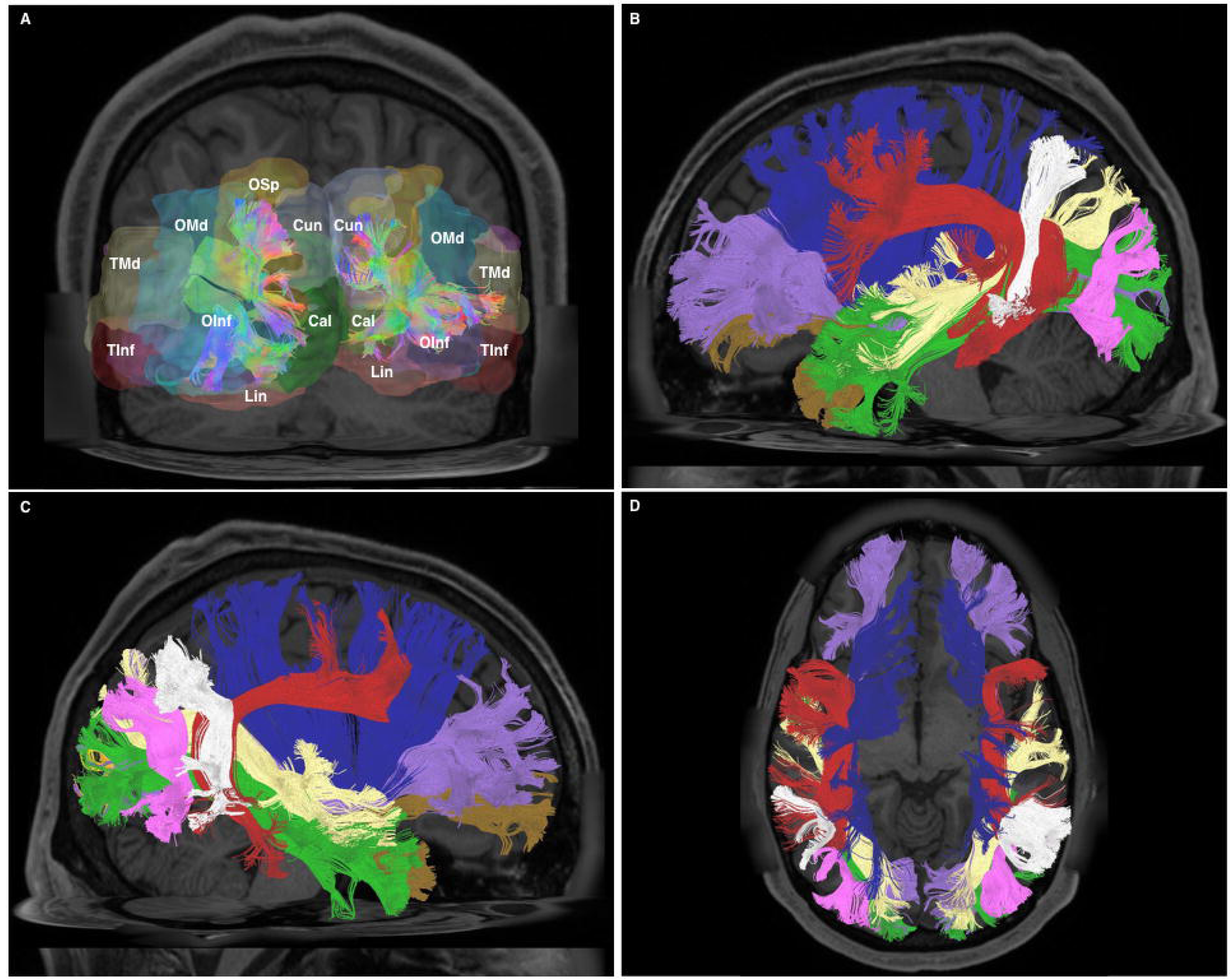
7A - Posterior-coronal view demonstrating occipito-temporal terminations of all divisions of the ILF. OSp - superior occipital gyrus, OMd - middle occipital gyrus, Olnf - inferior occipital gyrus, Lin - lingual gyrus, Cun - cuneus, Cal - calcarine gyrus, TMd - middle temporal gyrus, Tlnf - inferior temporal gyrus. 7B - Left-hemispheric sagittal-view representation of relational white matter tracts. Tracts are colored separately. lFOF - purple, Claustrum - blue, AF - red, PAT-white, MdLF - yellow, ILF - green, UF - brown, VOF - pink. 7C - Right-hemispheric sagittal-view representation of relational white matter tracts. Tracts are colored separately. IFOF - purple, Claustrum - blue, AF - red, PAT-white, MdLF - yellow, ILF - green, UF - brown, VOF - pink. 7D - Superior axial-view representation of bi-hemispheric relational white matter tracts. Tracts are colored separately. IFOF - purple, Claustrum - blue, AF - red, PAT-white, MdLF - yellow, ILF - green, UF - brown, VOF - pink.

### 3.3 Volumetry and Lateralization

Whole, merged ILF bundles had a mean volume of 19.1 ml (SD = 6.0), while right sided whole ILFs had a mean volume of 14.1 ml (SD = 5.4). This 5 ml difference was significant *(T =* 3.38, p = 0.001). For separated sub-fascicles, mean left dorsolateral ILF volume was significantly larger than right; at 9.8 ml vs 6.9 ml, respectively *(T =* 3.50, p = 0.001). Ventrolateral and ventromedial ILF bundles were similar sized; with mean left ventrolateral and ventromedial ILF volumes at 7.4 ml and 7.3 ml, respectively. These were larger than their right sided counterparts, which also displayed similar mean volumetry for ventrolateral and ventromedial of 5.5 ml and 5.4 ml, respectively. The ventrolateral ILF was not significantly volumetrically lateralized *(T* = 1.96, p = 0.055) but the ventromedial ILF was *(T* = 2.51, p = 0.015). For whole ILF bundles, LI was 0.31 demonstrating leftward asymmetry. Analyzed LIs of the dorsolateral, ventrolateral and ventromedial sub-fascicles were 0.35, 0.30 and 0.29, respectively.

### 3.4 White Matter Tract Relations as Revealed by Tractography

At its anterior aspect, The ILF shares terminations within the temporal pole with the ventral UF. It then travels posteriorly, lateral to the temporal horn. At its longitudinal-middle segment, temporal connections of the AF overlie it. Originating from the superior temporal gyri, the MdLF passes deep to the posterior limit of the Sylvian fissure and the AF, travelling obliquely to the dorsal occipital and superior parietal area, remaining dorsal to the ILF at all stages. At the temporo-occipital junction, the ILF travels posteriorly, joining the sagittal stratum inferiorly and remaining ventro-lateral to the IFOF. As such, the IFOF lies between the ILF and optic radiations, all of which traverse antero-posteriorly via the sagittal stratum to occipital terminations. As it passes over the basal temporal areas, i.e. the fusiform gyrus, the ILF is overlaid by the vertical occipital fasciculus, a short vertically oriented white matter tract originating from the basal occipito-temporal areas and connecting with the dorsolateral occipital gyri. At its posterior and occipital aspect, the ILF shares termination areas with the IFOF, MdLF, VOF and the optic radiations **(Figures 7B-D, Figure 8A-B).**

**Figure 8.**
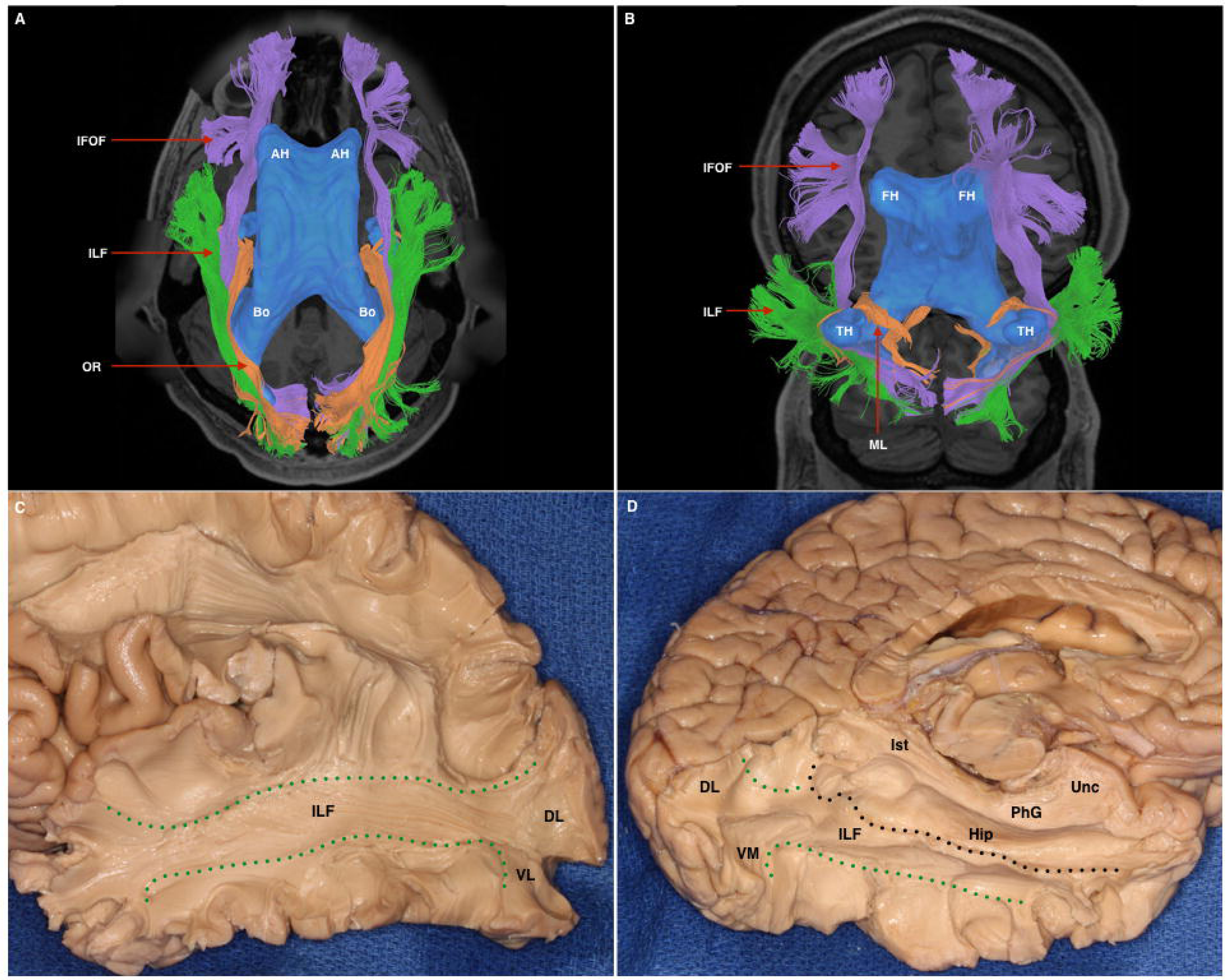
8A - Superior axial-view representation of tracts contributing to the sagittal stratum and their relation to the ventricular system. The ventricles are represented in blue as a 3-dimensional structure superimposed upon the axial T1 image. AH - anterior horn of the lateral ventricle, Bo - body of the lateral ventricle. lFOF - purple, ILF - green, OR - orange. 8B - Inferior para-axial representation of tracts contributing to the sagittal stratum and their relation to the ventricular system. The ventricles are represented in blue as a 3-dimensional structure superimposed upon the axial T1 image. AH - anterior horn of the lateral ventricle, TH - temporal horn. lFOF - purple, ILF - green, ML - Meyer’s loop, passing over the superior surface of the temporal horn. 8C - White matter dissection within a post-mortem specimen. Lateral-view. Cortex, U-fibers have been removed to expose the white matter of the ILF (demarcated by green broken lines) and the bifurcation of its two lateral (dorsolateral - DL; ventrolateral - VL) divisions. 8D - White matter dissection within a post-mortem specimen. View of infero-medial hemispheric surface. Superficial structures have been removed to expose the ILF (between broken green (inferior) and black (superior) lines. Also labelled are the ventromedial and dorsolateral posterior divisions. Superior to the broken black line is the hippocampus (Hip), the calcar avis (CA). Dorsal to these structures are the isthmus (lst), parahippocampal gyrus (PhG) and the uncus (Unc).

### 3.5 White Matter Dissection Results

From the lateral surface, the dissection started at the intersecting region of the parieto-occipito-temporal junctions, removing the U-fibers overlying the supramarginal and angular gyri. The vertical part of the SLF, also known as the parietal aslant tract, was exposed and removed to further expose the temporal AF, which was subsequently removed to expose underlying ILF fibers. ILF fibers were exposed along their antero-posterior course to the occipital pole. Towards the occipital pole, ascending fibers belonging to the vertical occipital fasciculus (VOF), were removed and then three posterior terminations corresponding to termination pattern were identified: dorsolateral, ventrolateral and ventromedial. From an inferior aspect U-fibers from the fusiform gyrus were gently removed to expose longitudinal ILF fibers. Lateral and medial occipito-temporal sulci were opened, as well as the lingual and parahippocampal gyri. The ILF represented the most lateral and ventral tract among other fascicles within the sagittal stratum. Its bifurcation in the sagittal plane led to identification of two components: ventral and dorsal. No evidence of amygdalar connectivity was found. Dissection results reinforced the tractographic results: The ILF lays ventro-laterally to the posterior sagittal stratum fibers, all coursing in parallel to terminate within the occipital lobe. The ILF ran lateral to the temporal horn and its floor. At the temporal pole, the ILF ended lateral to the UF. At the occipital pole, it was flattened between the VOF laterally, and the IFOF medially **(Figure 8C-D).**

## 4 Discussion

### 4.1 ILF Morphology, Subdivisions and Relations

We have demonstrated that GQI-tractography can reproduce bilateral ILF bundles according to previous descriptions. Our results confirm the existence of a robust, temporo-occipital fascicle originating from medial and lateral anterior temporal areas and terminating throughout the occipital lobe. We did not find any evidence of the ‘indirect’ U-fiber component of the ILF (Catani et al., 2003; Tusa and Ungerleider, 1985). Further, we were able to divide the whole bundles into smaller sub-fascicles for individual study. We used morphological termination patterns of the posterior ILFs to further subdivide them. For both left and right hemispheric ILFs, posterior arrangement was consistent over 30 subjects and within the HCP 842 atlas. Anterior ILF termination pattern was relatively more variable and did not demonstrate consistent morphology permitting subdivision. In terms of morphology and subdivision, our results are generally consistent with both older (Catani et al., 2002, 2003; Fernández-Miranda et al., 2008) and more recent (Latini, 2015; Latini et al., 2017) studies, all of which demonstrate a trifurcated posterior subdivision of the ILF. Though both Catani et al. (2003) and Latini et al. (2015, 2017) define a specific ‘cuneate’ branch of the dorsal subdivision, we opted not to use this as a specific criterion for subdivision due to its diminutive size and inconsistent representation across subjects (see also *ILF Connectivity* section). Regarding relations, we confirm that the ILF is the most superficial and ventral white matter fasciculus of the temporal lobe. Though cortico-cortical U-fibers may lie superficial to the robust ILF and indirectly connect temporal with occipital areas via short connections, we have found no tractographic evidence to suggest their anatomical affiliation with the ILF system.

### 4.2 Quantitative ILF Connectivity

We used a purely quantitative method of assessing the connectivity of the ILF and its sub-fasciculi. This represents a methodological evolution from our previous studies (Fernández-Miranda et al., 2015; Panesar et al., 2017; Wang et al., 2013, 2016), which all used qualitative correlations of fiber end-points within atlas regions. Not only does our quantitative technique give an objective measure of cortical termination, it also allows a metric analysis of the strength of connectivity between cortical areas. The need for quantitative termination analysis methods in tractographic studies has been identified (Girard et al., 2014) and will ultimately permit mapping of the connectome (Behrens and Sporns, 2012). Recently published quantitative analysis methods include statistical analysis of tract termination patterns derived from whole brain probabilistic DTI tractograms (Hau et al., 2016, 2017). Our use of connectograms (Yeh et al., 2018) aids understanding of connectivity strength by visually demonstrating bilateral ILF temporo-occipital connectivity in a topographically-correlated matrix.

From our connectivity analysis of individual sub-fasciculi, it was clear that all 3 sub-fascicles, and the merged ILF were leftward-lateralized in terms of connectivity. This is confirmed by an overall increased average connectivity strength on the left versus the right for individual sub-fascicles. Though our statistical tests did not show significant Cl lateralization for the dorsolateral or ventromedial ILFs, the ventrolateral ILF was significantly leftward-lateralized in terms of connectivity. Further, when taken as unseparated bundles, the left ILF demonstrated overall higher average Cl than the right. We postulate that the relatively small number of observations, and the large spread of Cl, contributed to the failure to demonstrate connective lateralization for dorsolateral and ventromedial ILFs. Moreover, though there were some fibers traversing between the cuneus, calcarine and lingual gyri, and the amygdala, these connections were negligible in relative Cl. We therefore question whether the ‘Li-Am’ bundle as proposed by Latini et al. (2015) is really a unique ILF sub-fasciculus, or rather could be attributed to tractographic artefacts.

### 4.3 ILF Volumetry

Our volumetric results, demonstrating significant leftward asymmetry of the whole ILF, and its individual constituents are in concordance with earlier DTI-derived results (Thiebaut de Schotten et al., 2011; Wakana et al., 2007), however it was in contrast to recent postulations by Latini et al. (2017), which demonstrated rightward volumetric asymmetry. Our results of leftward volumetric asymmetry can be attributed to overall increased leftward volume of all three ILF subcomponents, which all demonstrate LIs >0. Our results are in concordance with our previous study into IFOF structure, which also demonstrated leftward lateralization of this ventral tract. The ILF and IFOF may therefore be both structurally and functionally related.

### 4.4 Functional Postulations

Regarding function of individual subfascicles, the left dorsolateral ILF demonstrated strongest connectivity between the superior occipital gyrus and the two dorsal temporal gyri. The lateral occipital lobe, containing the superior, middle and inferior occipital gyri contains V3-V6 primary visual cortex, receiving direct input from the retina. The superior temporal gyrus has been implicated in emotional responses to visual facial stimuli (Domínguez-Borràs et al., 2009). The middle temporal gyrus has been implicated in both semantic access (Davey et al., 2015), distance judgement (Vandenberghe et al., 1996) and facial recognition (Haxby et al., 2000). The dorsolateral ILF is therefore strong candidate for a neural substrate sub-serving both semantic, spatial and facial recognition tasks. The left ventrolateral and ventromedial sub-fascicles connected with inferior and middle occipital gyri, cuneate, calcarine and lingual areas, respectively. Both had extensive terminations within the three lateral temporal gyri. Right sided ventromedial and ventrolateral ILFs demonstrated exclusive connectivity between lingual and the middle and superior temporal gyri. The cuneus is implicated in extrastriate visual processing (Allison et al., 1994). The calcarine cortex contains V1, and the lingual gyrus has been implicated in both facial recognition (Haxby et al., 2000) on the right and binocular spatial orientation bilaterally (Lee et al., 2001). Our results therefore implicate bilateral ventral components of the ILF to be implicated in facial recognition and would explain lesion-induced prosopagnosia (Bauer, 1984; Benson, 1974; Grossi et al., 2014; Michel et al., 1989; Takahashi et al., 1995).

Taken together, our results indicate strong leftward-lateralization of both connectivity and volume of the ILF, and its individual subcomponents. Within the ‘dorsal-ventral’ model of language functionality, prevailing opinion is of a leftward-lateralized ventral semantic system (Hickok and Poeppel, 2015). Our volumetric results therefore support a proposal of the ILFs role within the ventral semantic-stream. Further, due to its rich occipito-temporal connectivity, our results also support a dominant ILF role as an integrative system between the visual cortices and the relevant lateral temporal gyri, as evidenced by lesional studies (Bonilha et al., 2017). Recent theories (Dehaene and Cohen, 2011) have proposed the left-sided occipital visual wordform area may be a ‘functionally recycled’ neural substrate, originally evolved to subserve facial recognition. This theory is consistent with theories of ILF function, which implicate it in both semantic language and visual recognition roles. Our findings of ILF leftward-lateralization also reinforce this postulation.

### 4.5 Technical Considerations and Limitations

Our method was intended to generate whole ILFs without *a priori* regarding either connectivity or subdivision. Subsequent separation of the bundles did indeed reveal a tripartite ILF arrangement in accordance with the literature. Nevertheless, based upon our connectivity results, nomenclature for subdivided sub-fasciculi should not be used to indicate connectivity patterns, but instead only spatial orientation within the entire ILF bundle. This is exemplified by the right ventral ILF sub-fascicles, both of which demonstrate connectivity to both medial and lateral occipital cortices. Regarding connectivity overlaps, whether these occur due to true segregation between sub-fasciculi or whether these are due to tractographic artefacts remains to be determined. In an effort to reconcile this issue, and also proposed by Latini et al. (2017), we emphasize that both connectivity and volumetric results for whole, merged ILFs take precedent over results for individual sub-tract analyses. We also propose that connectivity-based segmentation be emphasized over morphological segmentation, as the former carries functional inferences as opposed to non-contextual spatial information of the latter.

Our current high-resolution GQI based imaging technique gives ability to accurately track close-proximity and crossing fiber systems to termination points within cortical mantle. The combination of this technique with quantitative connectivity analyses is the basis for connectogram construction and provides our most detailed and objective study to date. Further, when the pitfalls of DTI imaging are taken into consideration, the combination of tractography with objective connectivity analyses offers a superior solution for *in vivo* research studies. Moreover, though post-mortem fiber dissection has played a pivotal role in early anatomical studies and remains a valuable technique for comparison studies, its validity for connectivity analysis is very limited and widely surpassed by current tractographic methods.

## 5. Conclusions

We have successfully replicated a subdivided ILF bundle consistently over the range of 30 subjects. Though our results bear resemblance to previous postulations regarding the subdivision and gross connectivity of the ILF, we have offered a detailed picture of its bilateral connectivity patterns. The ILF is a connectively and volumetrically leftward-lateralized, ventral temporal bundle with rich occipito-temporal connectivity. Though tractography cannot give direct functional insight, our connectivity analysis taken in context with functional data pertaining to cortical functions offer an accurate and confirmatory insight into its postulated roles. We conclude by calling for increased adoption of GQI tractography and quantitative connectometry for anatomical studies.

## 6. Ethics Statement

This study was carried out in accordance with the recommendations and approval of the Institutional Review Board at the University of Pittsburgh. All subjects gave written informed consent in accordance with the Declaration of Helsinki.

## Abbreviations

NB: Left and Right hemispheric connections are demonstrated with a suffix _L or_R
Tlnf: inferior temporal gyrus
TMd: middle temporal gyrus
TSp: superior temporal gyrus
Fu: fusiform gyrus
ol nf: inferior occipital gyrus
OMd: middle occipital gyrus
OSp: superior occipital gyrus
Cal: calcarine gyrus
Cun: cuneus
Lin: lingual gyrus
Am: amygdala
Hipp: hippocampus

